# NeuroMark-HiFi: A Data-Driven Method for Detecting High-Spatial-Frequency Functional Brain Networks

**DOI:** 10.1101/2025.05.02.651877

**Authors:** N. Behzadfar, A. Iraji, V. Calhoun

## Abstract

**Objective:** The Traditional functional neuroimaging approaches typically focus on low-frequency spatial structures, potentially overlooking critical fine-scale connectivity disruptions associated with brain disorders

**Methods:** We introduce NeuroMark-HiFi, a fully automated algorithm designed to enhance the detection of high-spatial-frequency functional brain network patterns. NeuroMark-HiFi systematically preserves and analyzes fine-grained network variations by integrating reference-informed independent component analysis (ICA), 3D high-frequency spatial filtering, and a frequency-informed ICA decomposition to extract high-frequency functional components with greater precision.

**Results:** Simulation studies and mathematical evaluations demonstrate that NeuroMark-HiFi significantly improves sensitivity to both individual and group differences driven by small local shifts in spatial patterns of intrinsic connectivity networks (ICNs). Compared to traditional methods, NeuroMark-HiFi revealed additional group differences between individuals with schizophrenia (SZ) and healthy controls (HC), particularly in the visual, sensorimotor, frontal, temporal, and insular networks.

**Conclusion:** NeuroMark-HiFi successfully captures biologically meaningful alterations in spatial network patterns that conventional approaches may miss.

**Significance:** By improving sensitivity to subtle brain network alterations, NeuroMark-HiFi holds promise for early diagnosis, treatment monitoring, neurodevelopment studies, aging research, and multimodal biomarker discovery, advancing the goals of precision psychiatry and neuroscience.

## I. Introduction

DURING rest, the brain exhibits temporally coordinated activity patterns, known as resting-state networks (RSNs), which can be identified using functional magnetic resonance imaging (fMRI). These networks closely align with activation patterns observed during task-based fMRI.

Analyzing RSNs offers valuable insights into large-scale brain functional integration and has been widely used to investigate abnormalities linked to neurological and psychiatric disorders [1-5].

One of the most widely used methods for studying RSNs is independent component analysis (ICA), a multivariate technique that treats the brain as a set of spatially independent yet temporally coherent networks rather than isolated voxels. Unlike univariate approaches, which assess individual voxel signals independently, ICA leverages higher-order statistics to capture interactions between brain regions, enabling the extraction of functionally meaningful networks. This approach has significantly advanced the understanding of brain connectivity in many different contexts [6].

Although ICA and other traditional RSN analysis techniques have been instrumental in uncovering large-scale connectivity patterns, they typically prioritize global spatial features and may not fully capture fine-grained spatial variations. A broader gap in the field lies in the limited attention paid to the spatial frequency content of functional networks—information that could be critical for identifying subtle neural disruptions and early markers of brain disorders. Functional networks are not static or uniformly distributed across space; rather, they exhibit complex spatial organization that reflects underlying neurophysiological architecture.

Higher spatial frequencies, in particular, encode localized functional patterns and may be especially sensitive to early or subtle abnormalities in brain function. The spatial frequency composition of brain networks—reflecting the granularity of functional structures—has been shown to vary with age, gender, and mental illness [7], further highlighting the importance of incorporating multi-scale spatial information into RSN analysis. However, widely used methods such as ICA and seed-based correlation often underrepresent these finer spatial details, potentially overlooking important functional biomarkers.

Multidimensional frequency domain analysis techniques, such as the four-dimensional Fourier Transform (4D FFT), have been employed to investigate the spatial frequency composition of fMRI data [8]. These studies suggest that spatial frequency components contribute unique information about brain organization; however, they have not been explicitly designed to track functional network changes. This limitation presents a challenge, as small-scale shifts in functional networks can be difficult to detect without considering high-spatial-frequency components. For instance, variations in local connectivity due to neurodevelopmentaldisorders, psychiatric illnesses, or even normal aging may not be captured effectively through traditional spatial resolution approaches. This motivates the need for new analytical frameworks that preserve and analyze fine-scale spatial variations, ultimately improving sensitivity to network disruptions.

Beyond temporal analysis, spatial-frequency information in fMRI analysis has gained increasing attention. Studies suggest that high spatial frequency components in fMRI can provide valuable insights into fine-grained brain organization, revealing subtle connectivity structures that may not be apparent in conventional analyses. For example, empirical mode decomposition (EMD)-based decomposition methods have been employed to extract high spatial frequency features, which are linked to underlying neurophysiological sources and can enhance the precision of brain network characterization [9]. Additionally, ICA has been adapted to assess fine-grained functional structures by integrating spatial priors and high-frequency filtering techniques [10]. However, little work has focused specifically on the high spatial frequency information contained within extracted brain networks. This is particularly important because subtle network shifts—both across individuals and within groups—are likely to carry meaningful biological information.

Given ICA’s success in identifying RSNs at conventional spatial frequency ranges, extending multivariate methods to explore high spatial frequency representations is a logical progression. By integrating high-frequency spatial filtering into ICA-based analyses, we enhance sensitivity to subtle yet significant spatial variations in resting-state networks. This approach may not only more accurately capture typical variations but also improve the detection of subtle changes in the spatial patterns of networks and their alterations under different conditions.

In this study, we introduce NeuroMark-HiFi, a novel framework designed to improve the detection of high-spatial-frequency functional brain network patterns. Our method integrates spatially constrained ICA, 3D high-frequency spatial filtering, and a second ICA decomposition to extract fine-grained functional components with greater precision. We first outline the approach and discuss a mathematical approximation which illustrates how our approach can capture localized shifts in brain regions or networks which can be approximated by the spatial derivative. Next, to evaluate the sensitivity of our approach in detecting subtle spatial variations, we conducted a controlled simulation where spatially distinct components were systematically shifted between groups. Finally, we apply NeuroMark-HiFi to schizophrenia (SZ) data and observe a boost in sensitivity to group differences. Beyond SZ, NeuroMark-HiFi holds promise for advancing biomarker discovery, early diagnosis, and treatment monitoring in a variety of neurological and psychiatric disorders.

## II. METHODS

The proposed method consists of three key modules:

1. **Spatially Constrained ICA for Network Estimation** We begin with NeuroMark, a robust ICA-based framework for estimating brain functional network measures from fMRI data. NeuroMark enables reproducible identification of intrinsic connectivity networks (ICNs) across datasets, studies, and disorders [11]. Given fMRI data for a subject, denoted as *X_i_ E R^TxV^*, where *T* is the number of time points and *V* is the number of voxels, we extract *C_l_*ICNs per subject using spatially constrained ICA.
2. **Enhancement of High-Frequency Spatial Features** To highlight subtle spatial variations in brain networks, we apply a 3D high-pass spatial filter that suppresses low-frequency components while preserving high-frequency details. This ensures the maintenance of critical high-frequency spatial information. The filtered components retain C_l_ spatially refined ICNs per subject.
3. **Extraction of High-Frequency Spatial Patterns** The filtered networks from all subjects are concatenated and undergo a blind ICA (frequency-informed ICA) decomposition to extract high-frequency spatial patterns that exhibit covariation across subjects. This step isolates fine-scale variations that conventional ICA methods might overlook. Using frequency-informed ICA, we estimate subject-specific component loadings and extract C_2_ HiFi-components for each subject.

Fig.1 presents an overview of the NeuroMark-HiFi approach, which integrates spatially constrained ICA, high-frequency filtering, and high-frequency component extraction to enhance sensitivity to subtle spatial variations in brain networks.

**Fig. 1.**
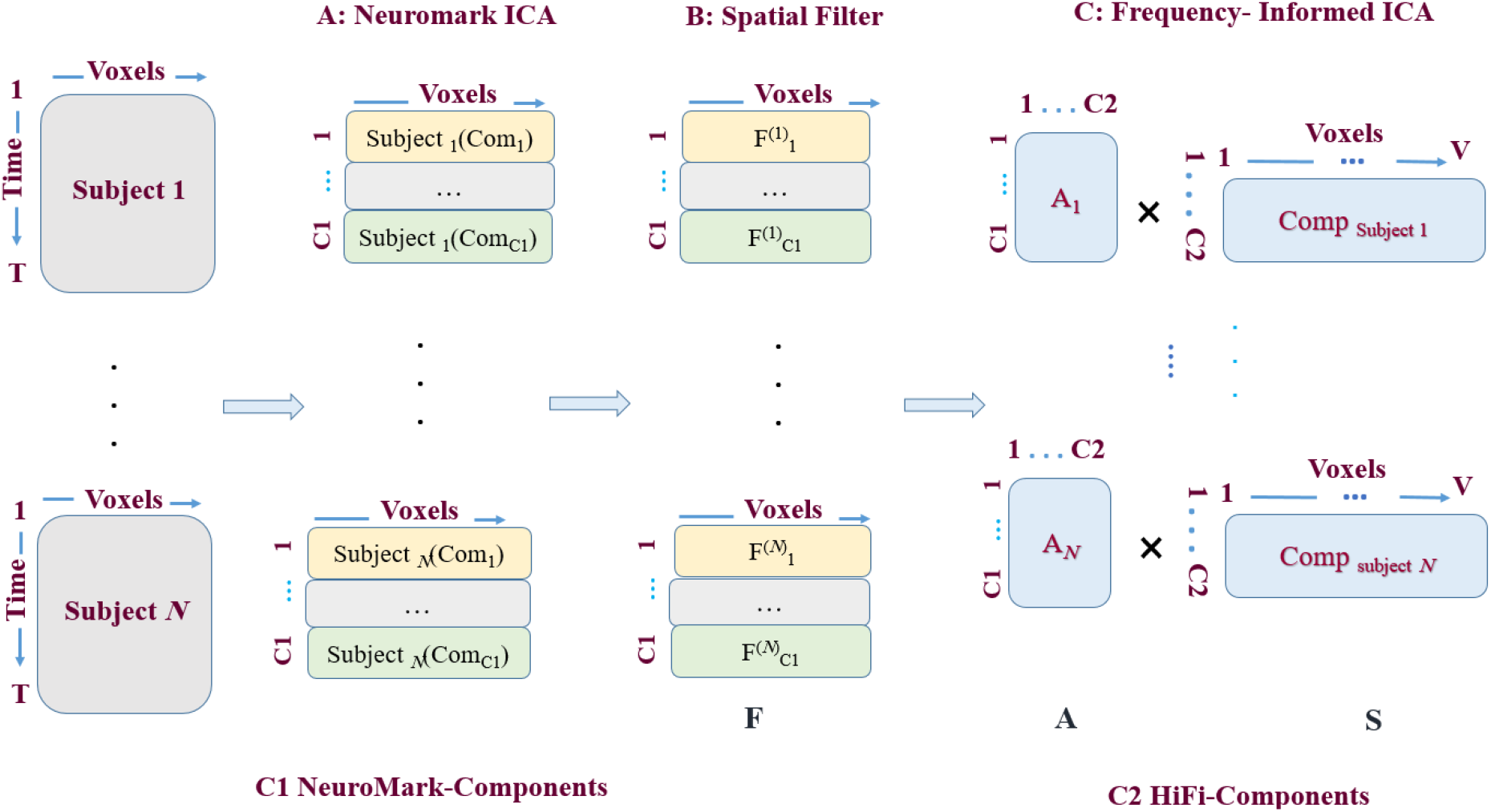
NeuroMark-HiFi Pipeline for Detecting Disease-Related Network Alterations. The pipeline consists of three steps: A) Spatially Constrained ICA extracts reproducible ICNs as templates from fMRI data; B) Spatial filtering enhances high-frequency features and suppresses background signals; and C) A frequency informed ICA is applied to the combined filtered networks across subjects to isolate coherent high-frequency components.

### Mathematical illustration

To illustrate the effectiveness of this approach, we first provide a mathematical description of how small spatial shifts in network components influence high-frequency spatial features. This is followed by a simulated experiment designed to demonstrate the method’s ability to detect subtle spatial shifts.

### Differences between two shifted blobs approximates the spatial derivative

Consider a brain network component represented as a 2D spatial image *l* (*x*.*y*) . with dimension 100 □100. We model two non-overlapping focal components, each with a full width at half max (FWHM) of 12 voxels. These components are represented as Gaussian blobs, centered at (*x_A_*.*y_A_*). for Source A and at (*x_B_*.*y_B_*) for Source *B* resulting in two sparse sources.

To introduce a subtle spatial difference between groups, we apply a small shift *δ*_y_ only to the blob in Source B in Group 2, representing source shifting between subjects, while the position of the blob for Group 1 remains unchanged, that is:

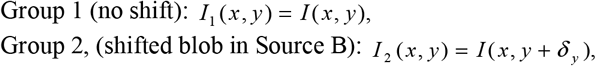

For small *δ_y_*, we approximate the difference using a first-order Taylor expansion:

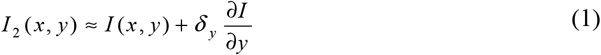

Thus, the difference between the two groups is:

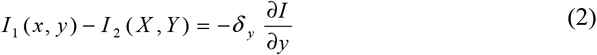

This implies that the spatial shift in the blob within Source B introduces a localized gradient-like difference in the image, proportional to the spatial derivative 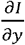. Next, we show that a simple high-pass filter approximates the spatial derivative.

### A high-pass filter approximates the spatial derivative

To enhance subtle spatial differences, we applied a high-pass filter in the frequency domain. A high-pass filter effectively removes/reduces low-frequency components while preserving/amplifying high-frequency details. This process is closely related to spatial differentiation, as differentiation in the spatial domain corresponds to multiplication by frequency components in Fourier space. For simplicity, we show this in 1D by considering the following discrete high-pass filter which is a common edge-detection filter (e.g., Sobel or Prewitt operator):

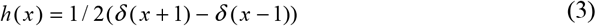

To apply this to a 1D spatial signal we convolve the signal *f*(*x*)with the filter *h*(*x*) :

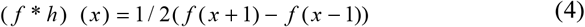

If we compare the above with the central difference approximation of the first derivative of the original signal *f* (*x*+ 1), we get:

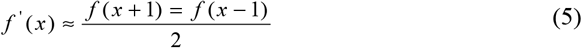

which exactly matches the convolution result in equation (4). We can also show this by taking the Fourier transform of *h*(*x*):

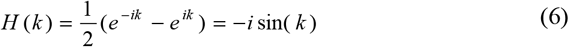

For small *k*, using sin k ≈ (*k*): *H*(*k*) ≈ *−ik*, which is exactly the Fourier transform of the first derivative.

### 2D simulation

To verify that **NeuroMark-HiFi** effectively captures high-frequency spatial group differences, we conduct a controlled simulation with the following steps:

1. **Generating Synthetic Brain Network Data** – We create a simulated 2D brain network composed of two Gaussian sources. Group 1 remains unchanged, while Group 2 experiences a slight **spatial shift in Source B**. Fig. 2.A (first row) illustrates the ground truth for two sources, A and B, as well as the ground truth for group differences 1 and 2. As shown, the difference between the two groups arises from a spatial shift in Source B.
2. **Application of High-Pass Filtering** – The simulated network images undergo **high-pass filtering** using the previously described filter to enhance localized variations.
3. **ICA and Spatial Components**-If we then apply ICA to the high-pass filtered images to identify independent spatial pattern, the observed data is modeled as a linear mixture of independent sources: **X= AS** where **X** is the matrix of observed high-pass filtered images (vectorized across subjects), **A** is the mixing matrix representing subject-specific loadings, and **S** is the matrix of independent spatial components.

**Fig. 2.**
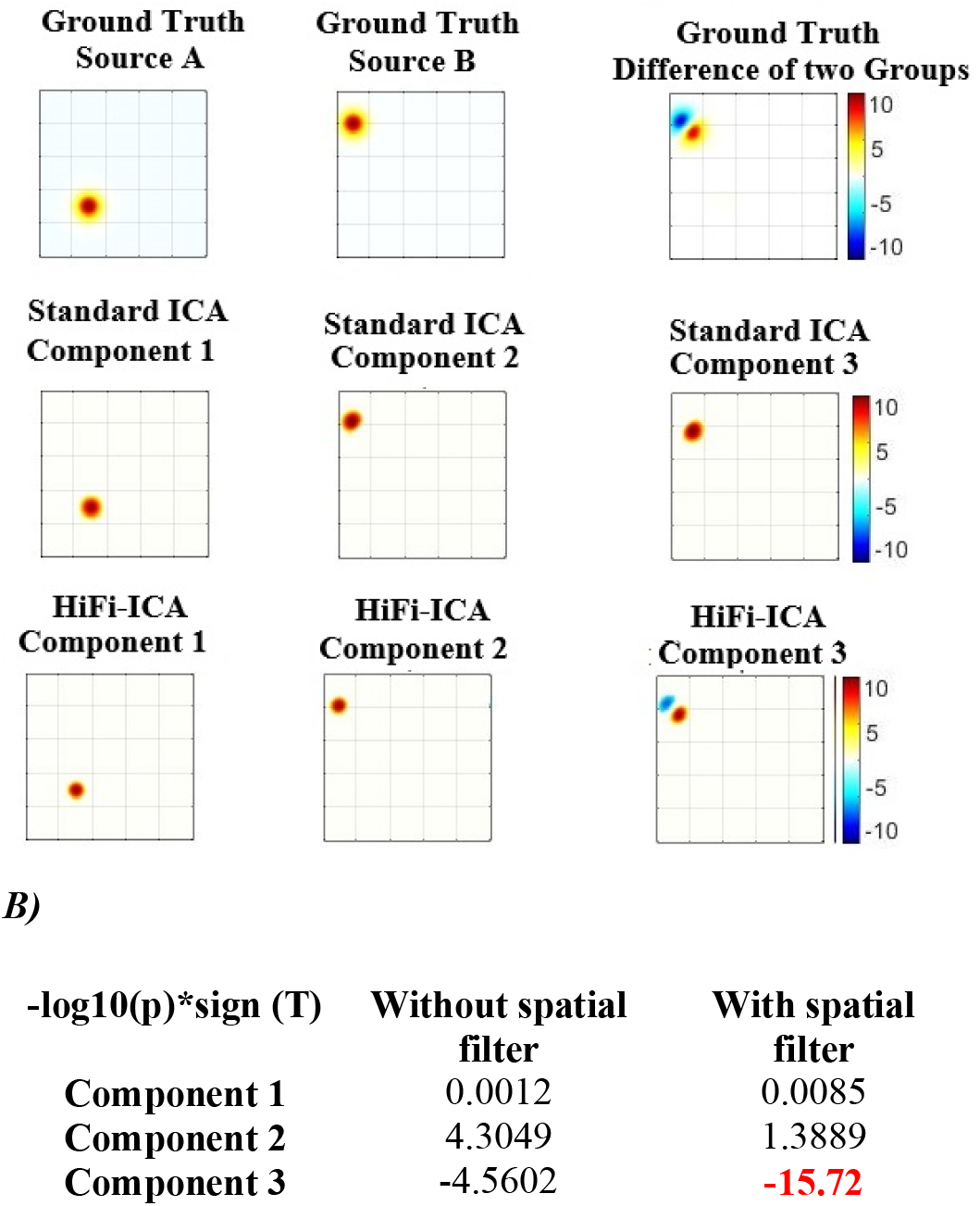
Sensitivity of NeuroMark-HiFi in Detecting Subtle Spatial Differences. (A) First row: Ground truth components without spatial shifts, serving as the baseline for comparison, including Source A, Source B, and the group difference map. Second and third rows: ICA components extracted using the standard ICA method (second row) and the NeuroMark-HiFi approach (third row), highlighting differences in detected spatial patterns. (B) Comparison of *−*log□□(p-values) for the three components across both methods. Notably, Component 3 in the NeuroMark-HiFi approach exhibits a highly significant difference between groups (T ∼ -15), whereas the variation for the standard ICA approach has been split across two components, with reduced effect (T ∼ 4), demonstrating the potential of NeuroMark-HiFi to enhance sensitivity in detecting subtle spatial variations.

**Fig. 3.**
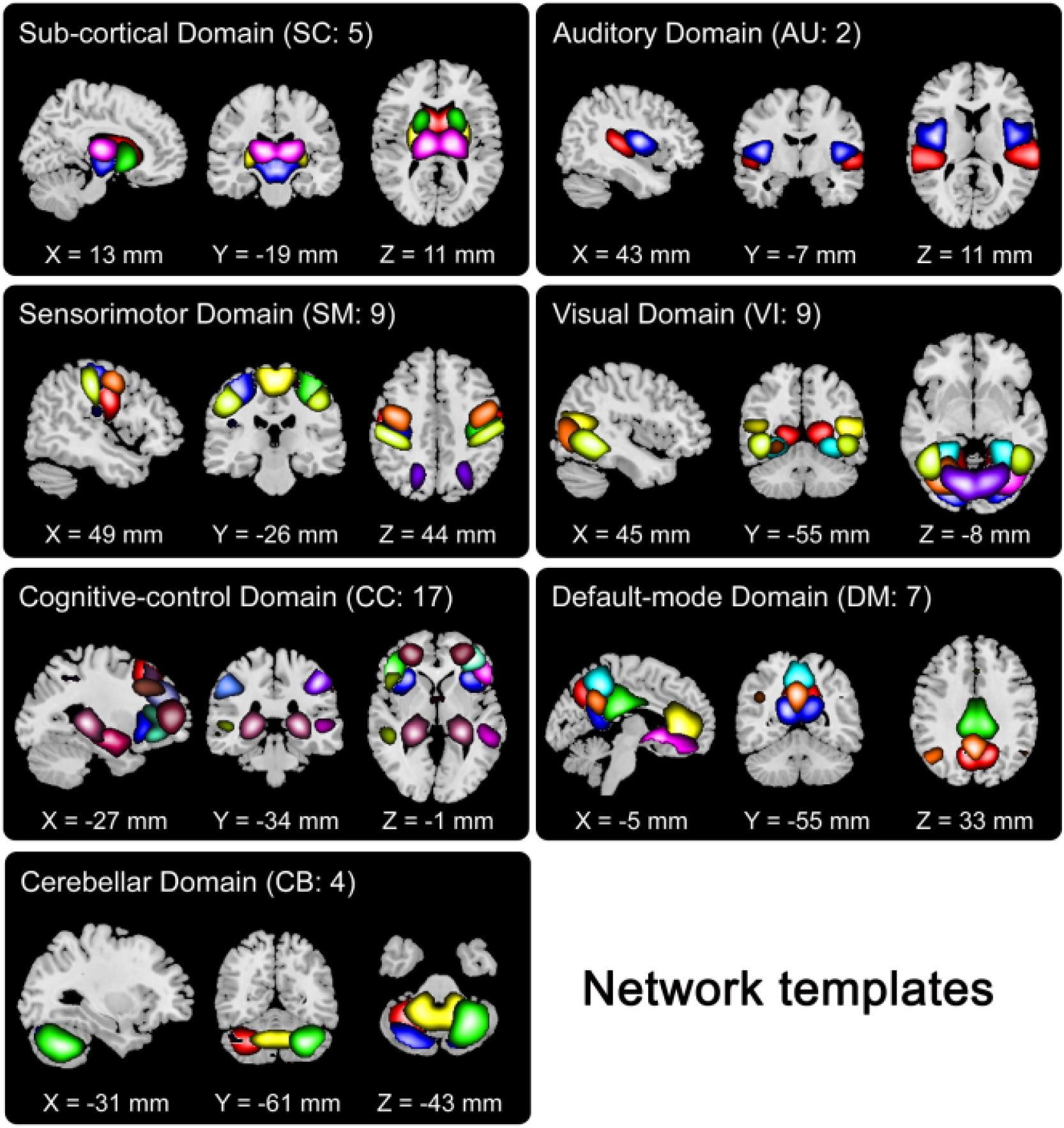
Visualization of NeuroMark network templates, which are divided into seven functional domains based on their anatomical and functional properties. In each subfigure, one color in the composite maps corresponds to an ICN.

Applying ICA estimates the unmixing matrix W leading to:**S=WX**.

Given *X_i_*as the fMRI data for the i-th subject, where *i ∈* {1,…, *N* } NeuroMark ICA is first applied to extract C1 ICNs for each subject. Following this, high-frequency filtering is applied, resulting in C1 filtered components per subject. Let 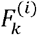 denote the k-th component after filtering for subject i.

Subsequently, frequency-informed ICA is performed on the concentrated filtered data, extracting C2 components. This process results in the estimation of C2 components across the dataset, which can be expressed as:

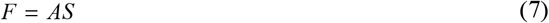

where:

- **F** is the aggregated matrix of filtered ICNs,
- **A** and **S** correspond to the mixing and source matrices, respectively.

For each subject, the mixing matrix consists of C1 values, indicating the expression of high-frequency-filtered ICNs. Through frequency-informed ICA, C2 informative components are extracted, resulting in C2 refined components per subject.

### Analysis of fMRI data

Next, we analyzed eyes-closed resting-state fMRI scans from 131 SZ patients and 141 healthy controls (HCs), obtained from the phase III dataset of the Functional Biomedical Informatics Research Network (fBIRN) [12]. The data acquisition and preprocessing steps are explained in [13]. In summary, echo planar imaging was employed to acquire volumes of BOLD data across seven sites, all utilizing 3T MRI scanners. The scans were performed with an echo planar imaging sequence FOV of 220 × 220 mm (64 × 64 matrix), TR = 2 s, TE = 30 ms, FA = 77°, 162 volumes, 32 sequential ascending axial slices of 4 mm thickness and 1 mm skip), and subjects kept their eyes closed throughout the scanning session. The preprocessing of the fBIRN dataset follows a standard pipeline using the Statistical Parametric Mapping (SPM12, http://www.fil.ion.ucl.ac.uk/spm/) software. These steps included rigid body motion correction to address head motion, slice-timing correction to account for timing differences in slice acquisition and warping into the standard Montreal Neurological Institute (MNI) space using an echo planar imaging (EPI) template. Additionally, the data was smoothed to a 6 mm full width at half maximum (FWHM) using AFNI’s BlurToFWHM algorithm. This smoothing step serves as a conservative denoising process during preprocessing and helps reduce scanner-specific variability in spatial smoothness, providing “smoothness equivalence” across data from multiple sites.

NeuroMark ICA was performed using the spatially constrained ICA framework implemented in the Group ICA of fMRI Toolbox (GIFT) in MATLAB (http://trendscenter.org/software/gift) [14, 15]. This approach enables the automated estimation and labeling of individual-subject connectivity features by incorporating spatial network priors derived from large independent samples [11]. The Moo-

Next, frequency-informed ICA was performed on the spatially filtered NeuroMark components using GIFT in MATLAB. Prior to ICA decomposition, subject-level principal component analysis (PCA) was applied for data normalization. The extracted principal components from each subject were then concatenated along the temporal dimension, followed by a second PCA step at the group level to reduce dimensionality while preserving the most variance in the data. These group-level components were then used as input for ICA to achieve group-level decomposition [16]. The Infomax algorithm, as implemented in GIFT, was employed for ICA. Subject-specific spatial maps and time series were obtained through a back-reconstruction approach, which reverses the PCA steps to project individual data onto the group-level reference, ensuring accurate subject-specific network estimation [17, 18]. The model order for the frequency-informed ICA was determined using the elbow method applied to a PCA scree plot. Specifically, eigenvalues of the covariance matrix from the input data were computed and plotted, and a variance threshold of 95% was used to guide the selection. This analysis indicated that 30 components would optimally capture the majority of variance. Accordingly, we set the model order of the second-level group ICA to 30 and performed decomposition using this fixed number of components. At this stage, 30 (C2) independent components were extracted and used in the subsequent analysis.

Group differences were assessed using a general linear regression, and statistical comparisons were performed using functions available in MATLAB. To control for multiple comparisons and reduce the risk of false positives, we applied the Bonferroni correction. Specifically, the significance threshold was adjusted based on the number of comparisons, setting the corrected p-value threshold at 0.05 divided by the number of tests. This correction ensures rigorous control of the family-wise error rate and strengthens the robustness of the reported findings. ICAR algorithm, as implemented in GIFT, was used for ICA decomposition. For template matching and labeling, we utilized the *NeuroMark_fmri_1*.*0* template and associated labels. Using this method, we extracted 53 (C1) ICNs from the dataset, which serve as network templates in the NeuroMark pipeline.

Afterward, we applied a 3D high-pass spatial filter with a frequency cutoff radius of 7 (in voxel frequency units), implemented in the fourier domain. Given the voxel dimensions, this cutoff corresponds to a spatial frequency of approximately 0.031 mm^-1^, which lies just above the effective frequency cutoff imposed by the prior 6 mm FWHM Gaussian smoothing (*≈* 0.066 mm□^1^). We tested different cutoff values ranging from 3 to 7 and found that a cutoff of 7 best preserved fine-grained spatial details in the ICNs while effectively suppressing low-frequency information.

## III. RESULTS

### Simulation

We implemented a simplified simulation to illustrate how **NeuroMark-HiFi**, through the combination of high-pass spatial filtering and ICA, can enhance the detection of small-scale **spatial differences between groups**. Specifically, the method is designed to improve sensitivity to structured spatial alterations—such as small shifts in spatial maps—that may not be clearly detected by standard ICA alone. While our toy example focuses on a localized displacement, NeuroMark-HiFi is more broadly applicable for capturing group-level spatial variations that manifest in higher spatial frequency ranges.

In the simulation, both groups share two underlying spatial sources (Source A and Source B), but Group 2 includes a small spatial shift in Source B. Components were extracted using both standard ICA and the proposed NeuroMark-HiFi method. As shown in Fig. 2.A (second and third rows), both methods identify Components 1 and 2 corresponding to Sources A and B, respectively. However, a critical distinction arises in Component 3: only NeuroMark-HiFi identifies a distinct bimodal component that aligns strongly with the group difference map and yields the lowest p-value—indicating statistically significant group separation.

This result demonstrates that by incorporating high-pass spatial filtering before ICA, NeuroMark-HiFi enhances the sensitivity of ICA to structured spatial variations between groups, especially subtle differences such as small shifts. Although the number of spatial sources remains two, the spatial shift between the groups introduces a new dimension of structured variation that is isolated by NeuroMark-HiFi into a separate component. In contrast, standard ICA fails to isolate this effect clearly and distributes the difference across multiple components with weaker group-level contrast.

Notably, the spatial filtering is not meant to enforce a frequency constraint within each component but rather serves to amplify group-level differences that reside in higher spatial frequency bands. Thus, NeuroMark-HiFi does not merely capture high-frequency content—it leverages it to enhance inter-group contrast when such differences occur at finer spatial scales. The observed p-value improvement (Fig 1.B) underscores this benefit. Overall, these results support the idea that NeuroMark-HiFi improves detection of subtle group-level spatial pattern variations, validating its potential for applications such as identifying disease-related alterations in brain networks.

By applying a 3D high-pass spatial filter, we suppress low-frequency information while preserving fine-grained spatial details in the ICNs. Figure 4 presents the mean of spatial maps of three specific Neuromark components (8, 19, and 43) across all subjects, shown before (left) and after (right) applying the spatial filter to the Neuromark components.

**Fig. 4.**
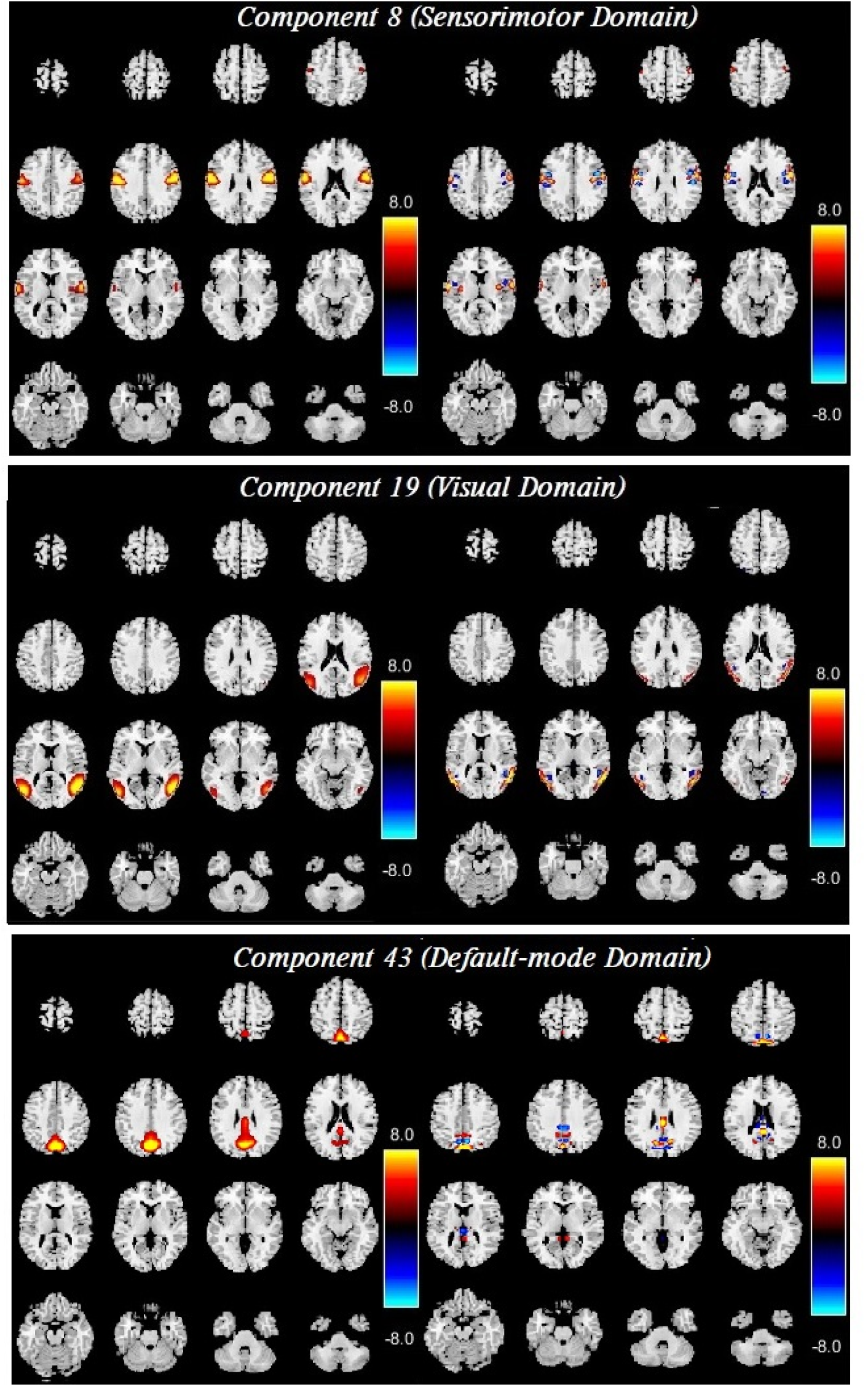
Spatial maps of three NeuroMark components before (left) and after (right) applying the high-pass spatial filter. A display threshold of 3 was applied, and the color bar represents the Z-score.

### Analysis of fMRI data

Using Spatially Constrained ICA, we extracted 53 NeuroMark components. Fig.3 illustrates the network templates employed for this extraction, categorizing them into seven functional domains based on their anatomical and functional characteristics. In each subfigure, distinct colors in the composite maps represent individual ICNs.

For each subject, we extracted 30 components (C2) from each of the 53 filtered ICNs (C1), resulting in a total of 1590 loading values (53 × 30). Among these, 14 components were identified as artifacts (e.g., edge or ventricular regions) based on visual inspection and spatial patterns, and were excluded from subsequent analysis. The remaining 16 components were further evaluated for group-level differences.

We identified components with significant group differences in their loading parameters using a general linear regression model, with diagnosis (SZ vs. HC) as the main predictor and age, sex, and scanning site included as covariates to control for potential confounding effects. To correct for multiple comparisons across 30 components, we applied the Bonferroni correction, setting a significance threshold at p < 0.05/30 *≈* 0.0016. This analysis yielded 12 components showing significant group differences, which we refer to as HiFi components due to their ability to capture high spatial frequency information across multiple networks.

These HiFi components are composed of contributions from multiple NeuroMark networks. In this context, the spatially filtered NeuroMark networks serve as spatial basis functions that combine in different ways to form each component. These components represent interpretable and high-resolution features, enhancing our ability to distinguish between groups.

The identified HiFi component spatial maps with p-values < 0.05 (corrected for multiple comparisons) are shown in Figures 5 and 6. In Figure 5, the most significant differences were observed in the frontal regions (HiFi components 4,11 and 20), sensorimotor regions (HiFi components 8 and 17), temporal regions (HiFi component 10) and visual regions (HiFi components 2 and 25). In these regions, SZ subjects exhibited significantly higher loading parameters than HC subjects, as indicated by positive and statistically significant regression coefficients.

**Fig. 5.**
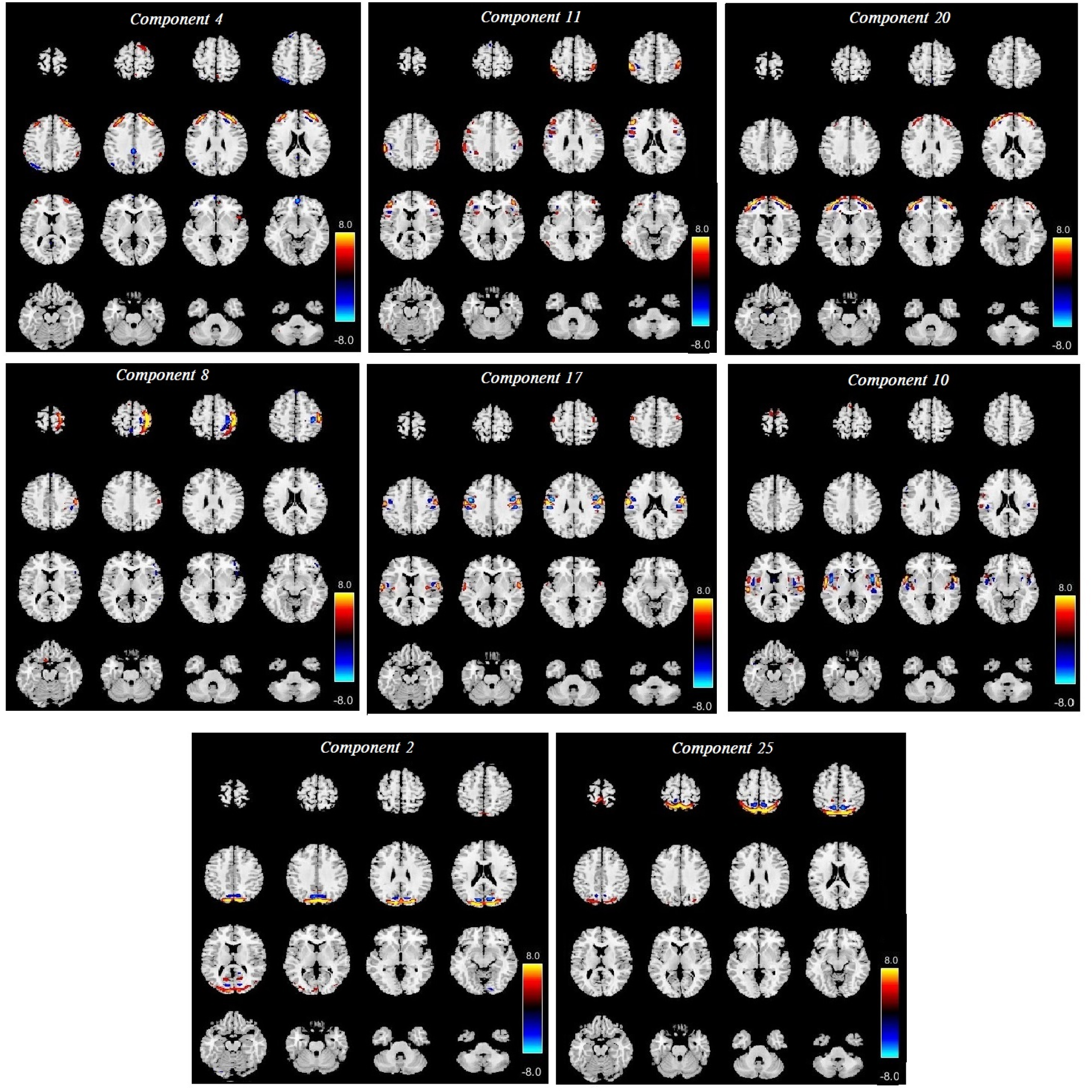
Spatial maps exhibiting significant group differences with SZ>HC are located in frontal regions (HiFi components 4,11 and 20), sensorimotor regions (HiFi components 8 and 17), temporal regions (HiFi component 10) and visual regions (HiFi components 2 and 25). In these regions, SZ subjects showed significantly higher loading parameters compared to HC, as indicated by positive and statistically significant regression coefficients. A display threshold of 3 was applied, and the color bar represents the Z-score.

**Fig. 6.**
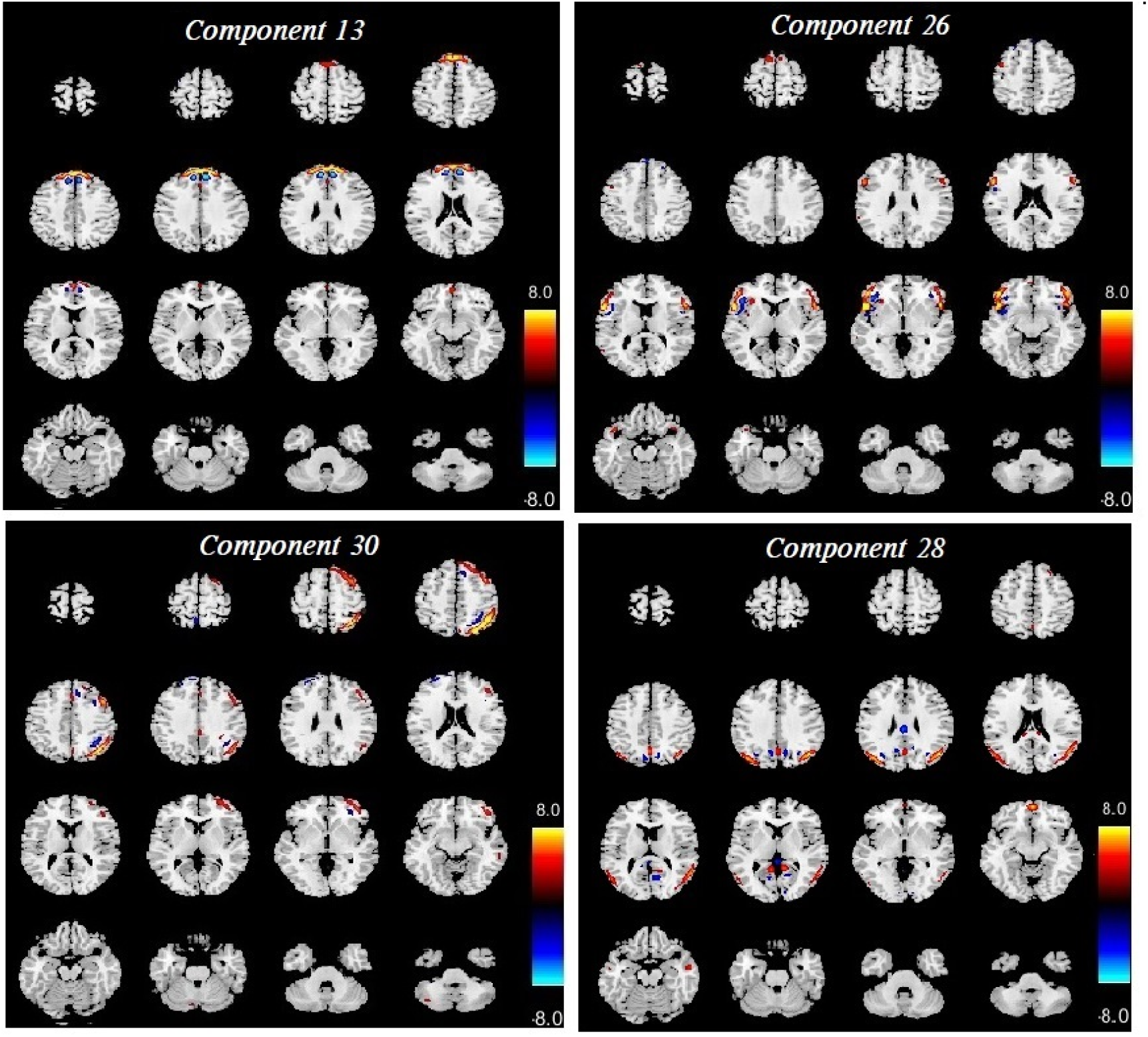
Spatial maps exhibiting significant group differences in HC>SZ are located in temporal region (HiFi component 13), insula/lateral frontal region (component 26), bilateral temporal region (HiFi component 28), and lateral frontoparietal region (HiFi component 30). In these HiFi components, SZ subjects exhibited significantly loading parameters compared to HC subjects, indicating reduced functional engagement in these regions. A display threshold of 3 was applied, and the color bar represents the Z-score.

In Fig. 6, the most significant differences were found in the temporal region (HiFi component 13), insula/lateral frontal region (component 26), bilateral temporal region (HiFi component 28), and lateral frontoparietal region (HiFi component 30). In contrast to the findings in Figure 5, SZ subjects exhibited significantly lower loading parameters than HC subjects, as indicated by negative and statistically significant regression coefficients

## IV. DISCUSSION

Our study introduces NeuroMark-HiFi, a novel framework designed to enhance the detection of fine-scale functional connectivity changes in the brain by leveraging high-spatial-frequency information. Traditional ICA-based methods primarily focus on low-frequency spatial structures, potentially overlooking subtle but biologically meaningful network alterations. By integrating spatially constrained ICA with high-pass spatial filtering and a second ICA decomposition, our approach isolates high-frequency functional components, improving sensitivity to small-scale spatial shifts.

We present a simple mathematical analysis to show that localized shifts in brain network components manifest as high-frequency spatial variations, which can be effectively captured using a high-pass filter. Simulations further validate that NeuroMark-HiFi significantly improves sensitivity to subtle spatial differences between groups, reinforcing its potential for detecting fine-grained neurophysiological changes.

Applying NeuroMark-HiFi to SZ data revealed enhanced sensitivity to group differences that were missed by conventional methods. Specifically, we observed meaningful connectivity alterations in key resting-state networks, including the visual, sensorimotor, frontal, temporal, and insular regions—areas consistently implicated in schizophrenia. Prior studies have shown that disruptions in the temporal and insular networks are associated with auditory hallucinations, which are among the core positive symptoms of schizophrenia [19, 20] . Frontal network abnormalities, particularly within the dorsolateral prefrontal cortex, have been widely linked to executive dysfunction, affecting working memory and cognitive flexibility [21]. Alterations in the sensorimotor network have been related to impaired motor coordination and abnormal movement patterns commonly observed in patients [22], while disruptions in the visual cortex have been associated with visual perceptual abnormalities, including illusions and distortions reported in schizophrenia [23]. These findings indicate that the spatial differences identified by NeuroMark-HiFi not only improve statistical sensitivity but also capture biologically plausible alterations consistent with the known pathophysiology of the disorder. By focusing on high-spatial-frequency components, NeuroMark-HiFi enables detection of fine-grained connectivity changes that may serve as valuable biomarkers for early diagnosis and disease monitoring.

Conventional ICA-based approaches are effective for identifying large-scale network disruptions but often fail to capture finer spatial variations that might be crucial in certain disorders. NeuroMark-HiFi addresses this limitation by explicitly focusing on high spatial frequencies, allowing for improved characterization of functional networks. Our findings align with recent studies which suggest that high spatial frequency components in fMRI provide valuable insights into fine-grained brain organization, revealing subtle connectivity structures that may not be apparent in conventional analyses. For example, EMD-based decomposition methods have been employed to extract high spatial frequency features, which are linked to underlying neurophysiological sources and can enhance the precision of brain network characterization [9]. This highlights the importance of leveraging high-spatial-frequency components to gain a more detailed understanding of functional connectivity patterns.

Our findings demonstrate that several components are significantly associated with mental illness. Specifically, the regression analysis revealed notable group differences in component loading parameters, highlighting how brain connectivity patterns differ in individuals with schizophrenia (SZ) across different spatial frequency bands.

Our work shows that components have significant relationships with mental illness. Specifically, the regression analysis revealed significant group differences in components, highlighting how brain connectivity varies in SZ across different frequency bands. The most striking variations were observed in regions such as the cerebellum network, visual network, cognitive control network, DMN, and sensorimotor network. These findings align with previous research indicating that these areas are critically involved in the neuropathology of SZ. For instance, altered connectivity in the cerebellum and cingulate cortex reinforces the idea that SZ involves disruptions in both motor coordination and cognitive control networks [24, 25] . Similarly, the observed differences in the temporal lobes and basal ganglia further support their established roles in auditory processing and motor function, both of which are often impaired in SZ [26, 27] . Additionally, patients with SZ have been reported to show transient decreases in network coupling within the visual and auditory networks, along with increased variability in intradomain connectivity[5]. There is also evidence of transient reductions in functional activity and connectivity, particularly in the subcortical domain, suggesting fluctuating network dynamics in SZ [28].

Our findings are consistent with, but extend, those of Garrity et al. [29], who reported that SZ is associated with altered temporal frequency and spatial location of the DMN. Similarly, Calhoun et al. [30] demonstrated that SZ patients show changes in posterior default mode regions, which is corroborated by our results in **Figures 5 and 6**, where the most significant changes were observed in these areas. Moreover, **Fryer et al. [31]** identified differences between SZ patients and healthy controls in regions such as the **posterior cortex, occipital lobe, and cerebellar lobes**. These observations have been further supported by other studies [5, 28, 32-37].

Importantly, the results in the current study go well beyond this related work as no study has focused on network-specific spatial-frequency differences in schizophrenia. The improved sensitivity of NeuroMark-HiFi suggests that conventional ICA-based approaches may overlook critical fine-scale network alterations that contribute to the pathology of SZ. By incorporating high-spatial-frequency filtering, our framework enhances the detection of subtle functional connectivity changes, providing a more comprehensive and nuanced view of brain network dysfunction.

## V. LIMITATION

While NeuroMark-HiFi enhances sensitivity to fine-scale connectivity changes, some limitations should be noted. First, while we focus on a basic high-pass filter, the optimal cutoff frequency for high-pass filtering needs further exploration, as it may vary across datasets and clinical conditions, and possibly across x, y and z directions. Second, the second ICA decomposition introduces an additional computational step and also requires manual inspection to label artifacts and networks, which may increase processing time and complexity. Future work should explore optimization strategies, such as machine learning-based parameter tuning, and automated labelling, to improve efficiency. In particular, approaches like those proposed by Salman et al. [38] for automatic labeling and ordering of brain activity/component maps could be leveraged to reduce manual intervention and streamline processing.

Additionally, while our framework successfully identifies fine-scale alterations, the biological interpretation of high-spatial-frequency components remains an open question. Future studies should investigate the neurophysiological basis of these components perhaps in the context of task-based studies or brain stimulation or using complementary modalities such as EEG or MEG.

## VI. CONCLUSTION AND FUTURE WORK

This study presents NeuroMark-HiFi, a novel ICA-based framework that enhances the detection of fine-scale functional connectivity changes in fMRI data by leveraging high-spatial-frequency information. Our mathematical analysis, simulations, and real-world application to SZ data collectively demonstrate that high-frequency spatial components contain meaningful neurophysiological information that is often missed by traditional methods.

NeuroMark-HiFi not only improves sensitivity to individual and group differences but also highlights the potential importance of high-frequency network components in understanding neuropsychiatric disorders. These findings suggest that high-spatial-frequency features could serve as valuable biomarkers for mental illness, facilitating earlier diagnosis, better treatment monitoring, and improved precision psychiatry approaches. Beyond SZ, the ability of NeuroMark-HiFi to detect subtle network alterations suggests its potential for application in a wide range of neurological and psychiatric conditions. Disorders such as Alzheimer’s disease, autism spectrum disorder, and major depressive disorder may also involve fine-scale connectivity disruptions that conventional analyses often overlook. Future studies could explore its utility in these domains, as well as in broader applications such as brain development and aging. Additionally, future work will focus on refining the technique, exploring its clinical applications, and integrating it with multimodal neuroimaging data. NeuroMark-HiFi represents a significant step forward in functional neuroimaging, providing new avenues for understanding the brain’s intricate connectivity patterns with greater precision.

Moreover, an important future direction is to explore spatial dynamics by incorporating approaches such as spatial dynamic subspace analysis and spatial chronnectome modeling. These methods have demonstrated the ability to capture transient, overlapping spatial patterns and the dynamic interplay between functional segregation and integration across time, which are particularly relevant for understanding complex disorders like schizophrenia [4, 5]. Incorporating NeuroMark-HiFi into a spatiotemporal framework could uncover hidden transient features and state-dependent changes in spatial organization that are not captured by static approaches. This would further enhance our ability to detect fine-grained and time-varying neurophysiological signatures, opening new possibilities for real-time brain monitoring and personalized interventions.

